# Protein secondary structure prediction with context convolutional neural network

**DOI:** 10.1101/633172

**Authors:** Shiyang Long, Pu Tian

**Author notes:** Membership list can be found in the Acknowledgments section.

## Abstract

Protein secondary structure (SS) prediction is important for studying protein structure and function. Both traditional machine learning methods and deep learning neural networks have been utilized and great progress has been achieved in approaching the theoretical limit. Convolutional and recurrent neural networks are two major types of deep leaning architectures with comparable prediction accuracy but different training procedures to achieve optimal performance. We are interested in seeking novel architectural style with competitive performance and in understanding performance of different architectures with similar training procedures.

**Results:** We constructed a context convolutional neural network (Contextnet) and compared its performance with popular models (e.g. convolutional neural network, recurrent neural network, conditional neural fields …) under similar training procedures on Jpred dataset. the Contextnet was proven to be highly competitive. Additionally, we retrained the network with the Cullpdb data set and compared with Jpred, ReportX and Spider3 server, the Contextnet was found to be more accurate on CASP13 dataset. Training procedures were found to have significant impact on the accuracy of the Contextnet.

**Availability:** The full source code and dataset have been uploaded at https://github.com/qzlshy/second_structure_model and https://github.com/qzlshy/ss_pssm_hhm.

## Introduction

Protein secondary structure is the local three dimensional (3D) ofganization of its peptide segments. In 1951 Pauling and Corey first proposed helical and sheet conformations for protein polypeptide backbone based on hydrogen bonding patterns [1], and three secondary structure states were defined accordingly. There were two regular secondary structure states: helix (H) and sheet (E), and one irregular secondary structure type: coil (C). In 1983 Sander [2] developed a secondary structure assignment method DSSP (Dictionary of Secondary Structure of Proteins), which classified secondary structure into eight states (H=*α*-helix, E=extended strand, B=residues in isolated *β*-bridge, G=3_10_-helix, I=*π*-helix, T=hydrogen bonded turn, S=bend and C= coil, the remaining). These eight states were often reduced to three states termed helix, sheet and coil respectively. The most widely used convention was that G, H and I were reduced to helix (H); B and E were reduced to sheet (E), and all other states were reduced to coil (C) [3–5].

Understanding protein function requires knowledge of their structures. Although many protein structures have been deposited in Protein Data Bank [6](www.wwpdb.org), far more sequences were known. Additionally, with present second generation and coming more efficient (and accurate) sequencing technologies, the gap between known sequences and known structures was expected to grow with accelerated speed. Considering the high cost of protein structure determination by experiments and rapid increase of available computational power, predicting protein structures using their sequence information computationally was a potentially practical solution. As the first step to predict 3D protein structures, protein secondary structure prediction has been studied for over sixty years [4]. Secondary structure prediction methods can roughly be divided into template-based methods [7–10] which using known protein structures as templates and template-free ones [3, 5, 11, 12]. Template-based methods usually have better performance, but do not work well with sequences that lack homologous templates. Template-free methods utilize sequence information alone. Q3 accuracy (i.e. with secondary structures labeled as H, E and C) based on template-free methods has been increased slowly from ¡70% to 82–84% [3, 4, 13, 14], gradually approaching the theoretical limit (88–90%) [13]. Three major factors that decide prediction accuracy were input features, predicting methods(algorithms) and training dataset. Input features, as the source of the information, have critical impact on accuracy. Utilizing multiple sequence alignment of homologous sequences rather than a single sequence has long been recognized as a way to improve prediction accuracy. The PSSM (position-specific scoring matrices) calculated by PSI-BLAST (Position Specific Iterative - Basic Local Alignment Search Tool) [15] has been widely utilized, which contributed significantly to the improvement of the Q3 accuracy to over 80% on benchmark datasets [4]. Compared with results from predicting using the sequence information only, Q3 accuracy has been increased by about %10 [16]. Some other input features, such as physio-chemical properties of amino acids and protein profiles generated using HHBlits, have been used in some secondary structure prediction models recently [5, 11]. Dataset used for training secondary structure prediction model has grown to several thousand sequences [3, 5, 11]. The sequence identity within each of these dataset was usually smaller than 25% or 30%.

Many different algorithms have been used to predict protein secondary structure in previous investigations, such as hidden markov models [17] and support vector machines [7, 18]. In recent studies, neural networks were widely used. DeepCNF (Deep Convolutional Neural Fields) [3] was an integration of CRF (Conditional Random Fields) and shallow neural networks, DeepCNF improved accuracy over several methods including SPINE-X, PSIPRED and Jpred by 1–4% on some data sets. LSTM (Long Short-Term Memory) bidirectional recurrent neural network [11] was developed to predict protein secondary structure states, backbone angles, contact numbers and solvent accessibility. MUFOLD-SS [5] used inception-inside-inception network to predict secondary structure states. 2C-BRNNs (two-dimensional convolutional bidirectional recurrent neural network) [12] was used to improve the accuracy of 8-state secondary structure prediction.

Present focus of protein secondary structure prediction studies was mainly on accuracy of secondary structure state, a very coarse classification. Potential value of these studies would only be fully embodied when combined with 3D structural prediction and design. It might well be that information representations learned in non-output layers can provide additional assistance for later stage structural studies. Therefore, searching for novel architecture flavors is meaningful by providing potentially unique and useful representations even if no significant Q3 accuracy improvement was achieved. We plan to investigate the value of non-output layer representations of various typical neural network architectures for secondary structure prediction in future work. In this work, we constructed a ContextNet (context convolutional neural network) and obtained higher accuracy than a few typical LSTM and CNN based architectures on Jpred and some other frequently used test datasets.

## Materials and methods

### Dataset and hardware

Five datasets were utilized in this study. Jpred dataset [19] and CB513 [20] dataset were downloaded from Jpred server (http://www.compbio.dundee.ac.uk/jpred/about.shtml). Jpred dataset contained a 1348-sequence training set and a 149-sequence test set, these sequences were selected representatives from SCOP superfamilies rather than constructed with a simple sequence identity cutoff. All the test and training protein sequences belong to different superfamilies. CB513 contains 513 non-redundant sequences, all sequences in which had been compared pairwise, and were non redundant to a 5SD cut-off [20].

CASP12 and CASP13 were downloaded from Protein Structure Prediction Center (http://predictioncenter.org/), and the target structures were used.

Cullpdb [21] dataset was downloaded from dunbrack lab (http://dunbrack.fccc.edu/PISCES.php). Cullpdb dataset was generated on 2018.11.26 (the percentage identity cutoff was 25%, the resolution cutoff was 2.0 angstroms and the R-factor cutoff was 0.25). We downloaded from PDB (protein data bank, www.wwpdb.org) the 9311 chains in the Cullpdb list. All the sequences in Cullpdb, CB513, CASP12 and CASP13 datasets are culled with CD-HIT server, sequence that had more than 25% identity to any sequences in the datasets was removed. So the sequence identity in all the datasets are less than 25%. We also removed sequences that failed in our described PSSM construction procedure (see below). Finally, we have 8601 sequences in Cullpdb dataset, 261 sequences in CB513 dataset, 35 sequences in CASP12 dataset and 15 sequences in CASP13 dataset. The Cullpdb dataset was arbitrarily divided into a 8401-sequence Cullpdb training dataset and a 200-sequence test dataset.

All networks were trained with GPU (GTX 1080Ti).

### The context convolutional neural network

Both local interactions due to neighboring residues in primary sequence and various non-local interactions due to tertiary interactions and electrostatic interactions participate in deciding secondary structure state of each residue. Explicit tertiary structure input is not available for sequences that do not have corresponding 3D structure available. PSSM in principle convey relevant non-local interaction information, but not in explicit and easy-to-decode form. Both convolution and LSTM architecture have the ability to capture non-local interactions through either maxpooling or recurrent operations. However, maxpooling, while expands the receptive field of convolutional networks, reduces image size and results in loss of information. Recurrent operation makes training process rather computationally intensive when compared with convolutional networks. The context module was constructed to increase the accuracy of state-of-the-art semantic segmentation systems [22]. The module used dilated convolutions to systematically aggregate multiscale contextual information without losing resolution. Dilated convolution can effectively increase the receptive field in the kernel without increasing the model parameters or the amount of calculation. With the dilated convolution, non-local interactions may be captured by fewer convolution layers. We therefore, hoping to better capture non-local interactions without engendering expensive training computation, constructed a context convolutional neural network (Contextnet, See Fig 1). The first 8 layer convolution results were concatenated together and the features of different receptive field were mixed. The operation concatenates tensors along one dimension, here we put the channels of different layers togeter and the number of output channels is the sum of the first 8-layer channels. In 5-8 layers the dilated convolution were used, strides of the dilated convolutions in layers 5-8 were 2, 4, 8 and 16 respectively. The kernel size used were 3×1, the activation function was relu in hidden layers, and the activation function in the output layer was softmax. The loss function was cross entropy and the network was built with Tensorflow. Detailed code can be found on github.

**Fig 1.**
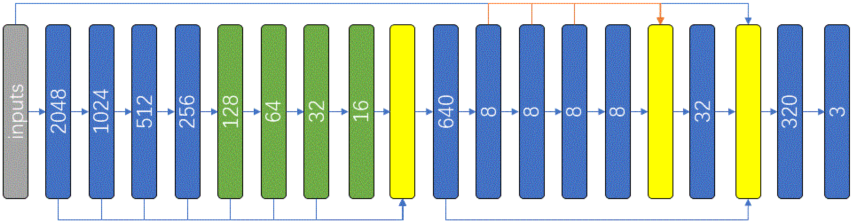
Context convolutional neural network architecture. The gray square represented input features, blue squares represented convolution operation, green squares represented dilated convolution operations and yellow squares represented concatenation operation. Numbers in squares were the channel number of the corresponding convolution operation.

### Input features and preprocessing

In Jpred dataset, PSSM were used as the only input features, with 20 elements to each residue. For Jpred dataset, both PSSM and labels were downloaded from Jpred server. For Cullpdb dataset, we ran PSI-BLAST with E-value threshold 0.001 and iteration number 3 to search UniRef90 [23] to generate PSSM. The UniRef90 database was downloaded from Jpred server. For Cullpdb dataset we also used physio-chemical properties [24] and HHBlits profiles [25] as input features. For a residue we had 57 features, 20 from PSSM, 30 from HHBlits profiles and 7 from physio-chemical properties. The HHBlits profiles were generated with uniprot20_2013_03 datasbase. We transformed the features to center it by subtracting the mean value of the training dataset, then scaled it by dividing by their standard deviation. The mean and standard deviation (obtained from the training set) were applied to the test set. Labels were obtained by first calculating 8-state DSSP labels and then reduced to 3-state, H (G, I, H), E (E, B) and C (others). The input features PSSM of each sequence were converted to shape (sequence length, 1, 20).

### Comparison with six other networks

In the first part of our performance comparison study, Adam (adaptive moment estimation) optimizer was used to train all the networks. For each network we manually chose an optimal learning rate among various tested values. The relu activation function was selected for all hidden layers and softmax was selected for the output layer, and with cross entropy selected as the loss function for all networks.

### Training strategies for the Contextnet

In performance comparison with servers, we trained the Contextnet with SGD (stochastic gradient descent) optimizer. To improve generalization capability of the Contextnet, a number of tricks were utilized. First we added white noise to input features by multiplying a random number sampled from a gaussian distribution with mean 1.0 and standard deviation 0.5 to each feature element; Second a random learning rate sampled from a uniform distribution in the range (0.02, 0.12) was utilized in each epoch of SGD optimization; Third a L2 regularization factor of 0.01 was utilized; Fourth, for each sequence half of the labels were randomly selected and masked (masked labels did not participate in backpropagation); Finally, two dropout layers were added (after the first and the third concatenation layers) in the network. The Q3 accuracy on corresponding test dataset was calculated every epoch during the training. The best test results from 20 epochs and corresponding model parameters were chosen as our final results. We did not use sequences that contain less than 20 residues when training with the Cullpdb dataset. The batch size was one sequence. We did not limit the size of convolutional network, the output feature numbers were the same as the length of the input sequence and each feature corresponds to a multi-class operation. So only one pass of convolution operation was performed on a sequence and results were obtained for all residues. The input feature shape was again (sequence length, 1, 20).

## Results

### Performance of the Contextnet and six other typical neural networks with the same simple training procedure

To evaluate the performance of the contextnet and to compare with other published networks, we first used Jpred dataset. To make the comparison relatively fair, all networks were given PSSM as the input features. The labels were 3-state secondary structure and were generated with DSSP calculations followed by a 8-state to 3-state reduction H (H), E (E, B), C (others). We trained the networks with Jpred training set and calculated the Q3 accuracy on test set every epoch. 10 to 20-epoch optimization was carried out for each network until apparent overfit was observed. Network parameter set that gave the highest test accuracy in epoches were chosen as the final result. We constructed seven different networks with Tensorflow. Some of networks were constructed according to previous studies (e.g. deepCNF, bidirectional recurrent neural networks and inception-inside-inception networks). No dropout and other special training methods (e.g. label masking, random noise addition, random learning rate etc.) were used in the training of all networks. It was important to note that the networks and training methods were not completely consistent with previous studies. Details were listed below:

1. Simple convolutional neural network:

A simple convolutional neural network with twelve hidden layers and one output layer was constructed. The kernel size was 3×1 and the number of channel was 256 in hidden layers. The SAME padding was utilized for each layer.

2. bidirectional LSTM neural networks:

The network was constructed with a bidirectional LSTM layer, two hidden layers and an output layer. The number of units in LSTM was 256. 1024 units were in the first and 512 units were in the second hidden layer.

3. Convolutional neural network with Conditional Neural Fields layer [3]:

The difference between Deep Convolutional Neural Fields (DeepCNF) and this implementation was that we used the relu activation function instead of sigmoid for hidden layers and we trained the network with Adam method instead of L-BFGS. Additionally, Wang trained the network layer by layer, but we trained the whole network directly. L2 regularization was not used here.

4. inception-inside-inception(Deep3I) [5]:

In our implementation no dropout layer was used and the input features were PSSM alone. In the original work physio-chemical properties of amino acids and HHBlits profiles were used as input features besides PSSM.

5. double bidirectional LSTM neural networks [11]:

Similar to 4), in contrast to the original work, the network was constructed without dropout layers. Physio-chemical properties of amino acids and HHBlits profiles were not used as input features.

6. Convolutional neural network with bidirectional LSTM layer:

A network with five convolution layers, followed by one bidirectional LSTM layer and one output layer was constructed. The kernel size was 3×1 and the channel number was 256 in convolution layers. The number of units in LSTM was 256.

7. Context convolutional neural network:

We constructed a context convolutional neural network (Contextnet). Concatenation and dilated convolution operations were utilized in the network. The detailed structure was described in the *Methods* section (see also Figure 1.).

We trained each network ten times. The average Q3 accuracy on test set and the standard deviation were shown in Table 1. The Contextnet obtained the highest accuracy at 83.96%

**Table 1.** Q3 accuracy of seven different secondary structure prediction networks on Jpred dataset.

|  | Accuracy(%) | Standard deviation (%) | Learning rate |
| --- | --- | --- | --- |
| Simple CNN | 82.85 | 0.26 | 0.0001 |
| BiLSTM | 83.26 | 0.12 | 0.001 |
| CNN+CRF | 82.16 | 0.26 | 0.0001 |
| Deep3I | 83.35 | 0.17 | 0.0001 |
| Double BiLSTM | 83.63 | 0.22 | 0.001 |
| CNN+BiLSTM | 83.68 | 0.21 | 0.0001 |
| Contextnet | 83.96 | 0.23 | 0.0001 |

These results by no means suggested that Contextnet was a superior secondary structure predictor to other tested networks as selection of training procedures and optimization details may change relative ranking of various networks. It was neither meaningful nor realistic to test all possible combinations of training methods. Nevertheless, the results strongly suggested that Contextnet was a competitive network.

### Training tricks and the ensemble method applied on the Contextnet

In order to improve the generalization capability of the network, we tested some training tricks on the Contextnet as described in the *Methods* section. With these tricks applied simultaneously, the Q3 accuracy of Contextnet was increased to 84.42% on the Jpred testset, and the variance also dropped as indicated by reduction of standard deviation from 0.23 to 0.10 (see Table 2)

**Table 2.** Improvement of Contextnet by training tricks and the ensemble method.

|  | Accuracy(%) | Standard deviation (%) |
| --- | --- | --- |
| Contextnet | 83.96 | 0.23 |
| Trained with tricks | 84.42 | 0.10 |
| Ensemble of Contextnet | 84.87 |  |

The performance can be further improved by the ensemble method. We retrained the Contextnet ten times and the ten resulting models were utilized to vote the final results. The Q3 accuracy increase to 84.87% as shown in Table 2.

### Comparison with Jpred, ReportX and Spider3 server on CB513, CASP12 and CASP13 datasets

Jpred [19], ReportX [26] and Spider3 [11] servers are widely known. To compare performance of our network with these servers, we retrained the Contextnet on the Cullpdb dataset, then tested the resulting model on CB513, CASP12 and CASP13 datasets. The input features used here were physio-chemical properties, HHBlits profiles and PSSM. We trained the model with 10 Cross-validation on Cullpdb training dataset and tested the model performance on Cullpdb test dataset. We further tested the model on CB513, CASP12 and CASP13 datasets to compare with Jpred, ReportX and Spider3 servers. The larger size (when compared with Jpred training set) of training datasets implied a potentially better evaluation of the performance for tested networks.

Jpred, ReportX and Spider3 server used different ways to reduce 8 secondary structure states to 3 states. Jpred reduced H to H, E and B to E, others to C. ReportX and Spider3 reduced H, I and G to H, E and B to E, others to C. Here we used the second reduced method. The results were listed in Table 3, 4 and the Ensemble of Contextnet provided higher accuracy.

**Table 3.** Q3 accuracy of Cullpdb 10 Cross-validation and test set.

|  | Mean accuracy of cv10 (%) | Mean accuracy of test (%) | Ensemble accuracy of test (%) |
| --- | --- | --- | --- |
| Contextnet | 84.90 | 84.51 | 85.32 |

**Table 4.** Q3 accuracy from Jpred server, DeepCNF server and the Contextnet.

|  | CB513 (%) | CASP12 (%) | CASP13 (%) |
| --- | --- | --- | --- |
| Jpred server | 80.11 | 78.49 | 80.01 |
| ReportX server | 82.34 | 80.84 | 81.13 |
| Spider3 server | 84.58 | 82.23 | 83.22 |
| Contextnet | 83.91 | 81.52 | 83.74 |
| Ensemble of Contextnet | 84.85 | 82.33 | 84.66 |

## Discussion

Performance of given networks may be improved by using combination of different hyperparameters, training methods (e.g. layer normalization, random noise, various regularization, dropouts etc.) and optimization methods [27]. We demonstrated that the same was true for the Contextnet when used to predict protein secondary structures. Significant impact of performance by application of various training tricks clearly illustrated that the free energy profile of neural network parameters have multiple local minima. Different training procedures, hyperparameters, initializations and optimization methods may take the training trajectory to various local minima with different generalization capability. Consistently, it was observed that all tested neural networks (with Jpred dataset) eventually overfit.

We made some effort to improve the Contextnet through various training tricks as described in the *Methods* section. Certainly, there might be more potential space for improvement that we simply do not have time to explore. The fact that we observed better performance for the Contextnet on nearly all tested datasets by no means suggested that the Contextnet was superior to other typical architectures, which may well be improved further in terms of Q3 accuracy if sufficient effort was made to explore combinations of training, optimization and hyperparameters. All published deep networks have the capability to extract non-local information and were sufficiently complex to overfit. Nonetheless, our contribution was that a new flavor of architecture that have competitive Q3 accuracy for protein secondary structure prediction was constructed and tested. As stated in the Introduction, diversity of network architecture might be of importance since intermediate representations learned might provide additional assistance to downstream 3D structural prediction and design. We were interested in exploring specifics of intermediate representations from various flavors of neural networks with competitive secondary structure prediction performance. Recurrent networks were significantly more expensive in training than convolutional networks while their performance as measured by Q3 accuracy were comparable. Nonetheless, we believed that recurrent networks were of great value as they can provide potentially unique useful information from intermediate representations.

## Conclusion

We constructed a context convolutional neural network to predict protein secondary structure state. In this network we used dilated convolutions to capture non-local interactions and used concatenation operation to mix multiscale contextual information. This network achieved competitive performance when compared with other published networks in our tests on seven data sets. In consistency with many neural network studies [27], we demonstrated the importance of training procedures in determining the generalization capability of the Contextnet. We believed that diverse architectures with competitive protein secondary structure prediction capability were potentially of great importance by providing different intermediate representations which might well be useful in downstream 3D structural studies. We plan to investigate specifics of learned intermediate representations from major network architectures used for protein secondary structure prediction.

## Acknowledgments

This work has been supported by the National Key Research and Development Program of China (2017YFB0702500), by the postdoctoral start up fund from Jilin University (801171020439), by National Natural Science Foundation of China (31270758), and by the Fundamental Research Funds for the Central Universities (451170301615).

## References

1. Pauling L, Corey RB, Branson HR. The structure of proteins; two hydrogen-bonded helical configurations of the polypeptide chain. Proceedings of the National Academy of Sciences of the United States of America. 1951;37(4):205–211.

2. Kabsch W, Sander C. Dictionary of protein secondary structure: pattern recognition of hydrogen-bonded and geometrical features. Biopolymers: Original Research on Biomolecules. 1983;22(12):2577–2637.

3. Wang S, Peng J, Ma J, Xu J. Protein secondary structure prediction using deep convolutional neural fields. Scientific reports. 2016;6:18962.

4. Yang Y, Gao J, Wang J, Heffernan R, Hanson J, Paliwal K, et al. Sixty-five years of the long march in protein secondary structure prediction: the final stretch? Briefings in bioinformatics. 2016;19(3):482–494.

5. Fang C, Shang Y, Xu D. MUFOLD-SS: New deep inception-inside-inception networks for protein secondary structure prediction. Proteins: Structure, Function, and Bioinformatics. 2018;86(5):592–598.

6. Berman HM, Westbrook J, Feng Z, Gilliland G, Bhat TN, Weissig H, et al. The protein data bank. Nucleic acids research. 2000;28(1):235–242.

7. Ward JJ, McGuffin LJ, Buxton BF, Jones DT. Secondary structure prediction with support vector machines. Bioinformatics. 2003;19(13):1650–1655.

8. Yi TM, Lander ES. Protein secondary structure prediction using nearest-neighbor methods. Journal of molecular biology. 1993;232(4):1117–1129.

9. Salamov AA, Solovyev VV. Prediction of protein secondary structure by combining nearest-neighbor algorithms and multiple sequence alignments; 1995.

10. Bondugula R, Duzlevski O, Xu D. Profiles and fuzzy k-nearest neighbor algorithm for protein secondary structure prediction. In: Proceedings of the 3rd Asia-Pacific Bioinformatics Conference. World Scientific; 2005. p. 85–94.

11. Heffernan R, Yang Y, Paliwal K, Zhou Y. Capturing non-local interactions by long short-term memory bidirectional recurrent neural networks for improving prediction of protein secondary structure, backbone angles, contact numbers and solvent accessibility. Bioinformatics. 2017;33(18):2842–2849.

12. Guo Y, Wang B, Li W, Yang B. Protein secondary structure prediction improved by recurrent neural networks integrated with two-dimensional convolutional neural networks. Journal of bioinformatics and computational biology. 2018;16(5):1850021–1850021.

13. Rost B. Protein secondary structure prediction continues to rise. Journal of structural biology. 2001;134(2-3):204–218.

14. Heffernan R, Paliwal K, Lyons J, Dehzangi A, Sharma A, Wang J, et al. Improving prediction of secondary structure, local backbone angles, and solvent accessible surface area of proteins by iterative deep learning. Scientific reports. 2015;5:11476.

15. Altschul SF, Madden TL, Schäffer AA, Zhang J, Zhang Z, Miller W, et al. Gapped BLAST and PSI-BLAST: a new generation of protein database search programs. Nucleic acids research. 1997;25(17):3389–3402.

16. Zvelebil MJ, Barton GJ, Taylor WR, Sternberg MJ. Prediction of protein secondary structure and active sites using the alignment of homologous sequences. Journal of Molecular Biology. 1987;195(4):957–961.

17. Karplus K, Barrett C, Hughey R. Hidden Markov models for detecting remote protein homologies. Bioinformatics (Oxford, England). 1998;14(10):846–856.

18. Cheng H, Sen TZ, Kloczkowski A, Margaritis D, Jernigan RL. Prediction of protein secondary structure by mining structural fragment database. Polymer. 2005;46(12):4314–4321.

19. Drozdetskiy A, Cole C, Procter J, Barton GJ. JPred4: a protein secondary structure prediction server. Nucleic acids research. 2015;43(W1):W389–W394.

20. Cuff JA, Barton GJ. Application of multiple sequence alignment profiles to improve protein secondary structure prediction. Proteins: Structure, Function, and Bioinformatics. 2000;40(3):502–511.

21. Wang G, Dunbrack Jr RL. PISCES: a protein sequence culling server. Bioinformatics. 2003;19(12):1589–1591.

22. Yu F, Koltun V. Multi-scale context aggregation by dilated convolutions. arXiv:151107122. 2015;.

23. Bairoch A, Apweiler R, Wu CH, Barker WC, Boeckmann B, Ferro S, et al. The universal protein resource (UniProt). Nucleic acids research. 2005;33(suppl_1):D154–D159.

24. Meiler J, Müller M, Zeidler A, Schmäschke F. Generation and evaluation of dimension-reduced amino acid parameter representations by artificial neural networks. Molecular modeling annual. 2001;7(9):360–369.

25. Remmert M, Biegert A, Hauser A, Söding J. HHblits: lightning-fast iterative protein sequence searching by HMM-HMM alignment. Nature methods. 2012;9(2):173.

26. Wang S, Li W, Liu S, Xu J. RaptorX-Property: a web server for protein structure property prediction. Nucleic acids research. 2016;44(W1):W430–W435.

27. Smith LN. A disciplined approach to neural network hyper-parameters: Part 1 – learning rate, batch size, momentum, and weight decay. arXiv:180309820. 2018;.

